# Maximizing the quality of NMR automatic metabolite profiling by a machine learning based prediction of signal parameters

**DOI:** 10.1101/466235

**Authors:** Daniel Cañueto, Miriam Navarro, Mónica Bulló, Xavier Correig, Nicolau Cañellas

## Abstract

The quality of automatic metabolite profiling in NMR datasets in complex matrices can be compromised by the multiple sources of variability in the samples. These sources cause uncertainty in the metabolite signal parameters and the presence of multiple low-intensity signals. Lineshape fitting approaches might produce suboptimal resolutions or distort the fitted signals to adapt them to the complex spectrum lineshape. As a result, tools tend to restrict their use to specific matrices and strict protocols to reduce this uncertainty. However, the analysis and modelling of the signal parameters collected during a first profiling iteration can further reduce the uncertainty by the generation of narrow and accurate predictions of the expected signal parameters. In this study, we show that, thanks to the predictions generated, better profiling quality indicators can be outputted and the performance of automatic profiling can be maximized. Thanks to the ability of our workflow to learn and model the sample properties, restrictions in the matrix or protocol and limitations of lineshape fitting approaches can be overcome.

## Introduction

Metabolomic studies characterize the low-molecular-weight components (<1 kDa) called metabolites in samples of biofluids or cell/tissue extracts.^1,2^ The quantification of the metabolite levels in nuclear magnetic resonance (NMR) spectra requires that the area below the metabolite signals to be quantified: this process is called metabolite profiling.^3,4^ This area can be quantified by area integration or signal deconvolution. In the case of 1D ^1^H-NMR spectra, three signal parameters need to be estimated to deconvolute a signal: intensity, chemical shift and half bandwidth.^3^ Once the combination of parameter values that fits the spectrum lineshape with lowest error has been estimated, the signal can be built and the area below the signal can be quantified. Several tools have recently appeared which can automatically estimate signal parameter values.^5–7^ These tools are usually based on optimization solvers (e.g., the Levenberg-Marquardt algorithm) which evaluate the search space shaped by the range of possible values of each parameter to find a minimum that represents the replication of the spectrum lineshape with the lowest fitting error.^8-9^

However, automatic approaches are compromised by the multiple sources of variability which can be observed in complex matrices (e.g., macromolecule-based baseline, chemical shift and half bandwidth variability –caused by pH, ionic strength or temperature fluctuations or signal overlap^10^) (Figure 1 (a)).These sources of variability oblige the ranges of the possible parameter values to be wider during lineshape fitting and, therefore, the presence of a wide range of local minima where the optimization algorithm can meet the completion criteria and, therefore, end into suboptimal resolutions (Figure 1 (b)).^11^ In addition, the possible presence of low-intensity signals adjacent to the ones of interest adds complexity to the spectrum lineshape. Consequently, optimization algorithms may not find the actual parameter values of the signals of interest but the ones which can best help replicate the complex lineshape. As a result of these challenges, automatic profiling tools sometimes provide wrong metabolite identifications (an important bottleneck in metabolomics^12^) and suboptimal quantifications. To reduce the generation of suboptimal fitting resolutions, several bioinformatic solutions can reduce the search space during optimization (e.g., the use of a chemical shape indicator [CSI], the simultaneous lineshape fitting of all the signals from the same metabolite or the modelling of chemical shifts using multiple sources of information, among others^13,14^). However, these strategies are dependent on prior information. Therefore, they cannot handle unidentified metabolites and might be not robust to small variations in the expected lineshape. For example, simultaneous lineshape fitting is prone to errors in case of chemical shift variability (additional examples of observed variations in the expected half bandwidth and in the expected intensity ratios of metabolite signals in a complex matrix-e.g., human urine-are available in Supporting Information). Consequently, to ensure optimal performance, some tools can only be used in specific matrices or require restrictive procedures in sample preparation and/or spectrum acquisition. These matrix- and protocol-based restrictions hinder the high-throughput potential of NMR or might mean the incorporation of false positives and negatives into the metabolomics literature when these restrictions are not strictly followed.^15,16^

**Figure 1.**
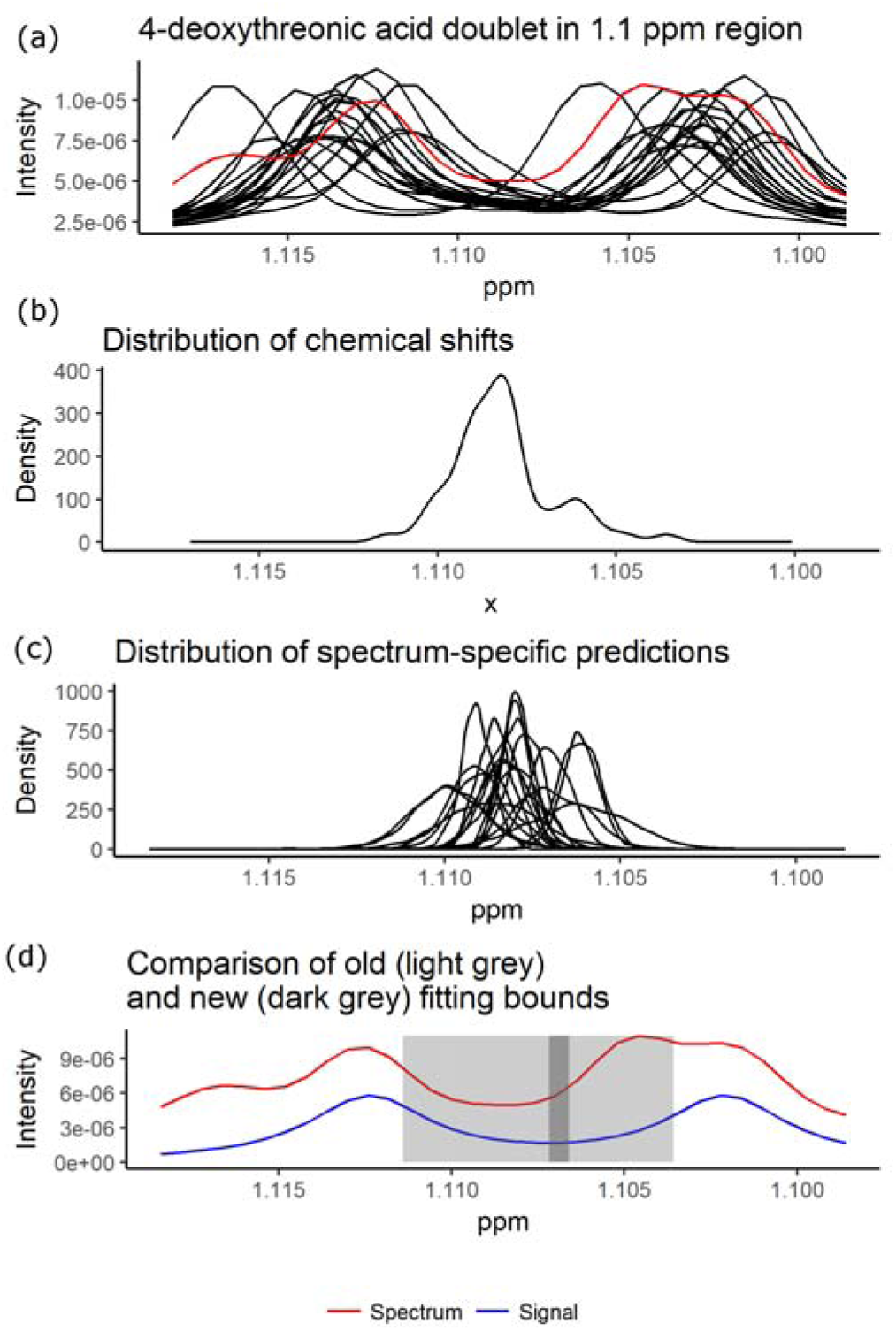
The signal parameter prediction pipeline enables narrow and accurate spectrum-specific ranges to be estimated and used during lineshape fitting. The figure shows a difficult signal fitting found with the 4-deoxythreonic acid signal in the urine dataset analysed in Supporting Information. The chemical shift variability present in this signal (a) forces lineshape fitting algorithms to consider a wide range of possible chemical shift values during the fitting (b). Excessive width can compromise the right assignment of the doublet centre when other signals appear adjacent to the signal to be fitted (d). The chemical shift prediction generates spectrum-specific chemical shift distributions of predictions (c). These distributions are very narrow and can help generate much narrower chemical shift ranges (d).

To maximize the quality of lineshape fitting during NMR automatic profiling, the ranges of possible parameter values selected during fitting must be as narrow as possible. Likewise, the estimation of these narrow ranges must be robust to the variable and complex properties of metabolomics study datasets in complex matrices. This combination of narrowness and accuracy can be achieved if the information about the sample properties necessary to narrow the ranges is collected from the same dataset during an initial profiling iteration. NMR signals mediated by atoms with similar chemical environments show similar reactivity to the fluctuations in the sample conditions. As a result, there is extensive multicollinearity in their half bandwidth and chemical shift values. This multicollinearity can be exploited to identify signals whose parameters do not behave as expected by this multicollinearity. In addition, accurate spectrum-specific predictions with prediction intervals (PIs) for each signal parameter can be estimated according to the information from the collinear signals. These PIs may be used to create very narrow and accurate value ranges to be used during lineshape fitting in a new profiling iteration. Likewise, the intensities of the signals from the same metabolite are perfectly collinear. Therefore, the expected intensity of each metabolite signal can also be predicted from the estimated intensities of the other metabolite signals. Consequently, the simultaneous lineshape fitting of all metabolite signals can be avoided. In contrast with other approaches, this prediction workflow is not dependent on prior matrix, protocol or metabolite information: this information is already encoded in the signal parameter values collected during the first profiling iteration. Therefore, it should be able to handle atypical or unidentified metabolites and be more robust to the sample-, matrix- or protocol-based complexities in the spectrum. In addition, the distance between the predicted parameter values and the collected parameter values can be quantified. The quantified distance may be a better profiling quality indicator than some of those in current use (e.g., fitting error) and help further minimise wrong annotations and suboptimal quantifications. To our knowledge, there has been no attempt to provide an open-source flexible automatic signal parameter prediction that maximizes the quality of the information provided by NMR profiling tools. In this study, we show how the proposed workflow helps maximize the quality of metabolite profiling in 1D ^1^H-NMR datasets.

## Methods

### Datasets

For this study, two datasets were analysed: a faecal extract dataset of 146 samples from a medical treatment study and a serum dataset of 212 samples from a nutritional intervention study.

In the faecal extract dataset, 70-100mg of dry faecal matter and 1000 ml of 0.05M PBS buffer in D2O were placed in a 2 ml Eppendorf tube. The sample was vortexed and sonicated until complete homogenization and the mixture was centrifuged (15000 rpm around 20000 g, 25 min, 4°C). For NMR measurement, 600 ml of the upper phase were placed into a 5mm NMR tube and ^1^H NMR spectra were recorded at 300 K on an Avance III 500 spectrometer (Bruker^®^, Germany) operating at a proton frequency of 500.20 MHz using a 5 mm PBBO gradient probe. One-dimensional ^1^H pulse experiments were carried out using the nuclear Overhauser effect spectroscopy. (NOESY) presaturation sequence to suppress the residual water peak, and the mixing time was set at 100 ms. The spectral width was 10 kHz (20 ppm), and a total of 256 transients were collected into 64 k data points for each 1H spectrum. After zero filling and exponential line broadening (0.5 Hz), spectra were Fourier transformed, manually phased and baseline-corrected using TopSpin software (version 3.2, Bruker). Bucketing (6e-04 ppm as bucket width), referencing to TSP at 0 ppm and probabilistic quotient normalization^17^ were performed through rDolphin.^18^

In the serum dataset, sample collection details are available in Hernández-Alonso, P. *et al*.^19^ For each sample, 300 μl aliquots were mixed with 300 μl of sodium phosphate buffer. Carr Purcell-Meiboom-Gill (CPMG) spectra, at 37°C and with presaturation to suppress the residual water peak, were acquired on a Bruker 600 MHz Spectrometer (Bruker Biospin, Rheinstetten, Germany) equipped with an Avance III console and a TCI CryoProbe Prodigy. CPMG data were pre-processed on the NMR console (TopSpin 3.2, Bruker Biospin, Rheinstetten, Germany) for zero filling, exponential line broadening (0.5 Hz) and phase correction. Bucketing (6e-04 ppm as bucket width) and referencing to the anomer of glucose at 5.233 ppm were performed through rDolphin.^18^

In addition, mass spectrometry (MS) profiling data was collected from both datasets. Complete details regarding the MS profiling workflow used in both datasets are available in Supporting Information.

### ^1^H-NMR metabolite profiling workflow

Automatic metabolic profiling was performed using the rDolphin R package^18^, an open source tool which as well collects the values of the signal parameters and exports them for analysis. rDolphin performs a lineshape fitting based profiling which adjusts spectral regions to a sum of Lorentzian signals, each one of which is characterized by three parameters: intensity, chemical shift and half bandwidth. The fitting process is performed using the Levenberg-Marquardt Nonlinear Least-Squares algorithm with lower and upper bounds provided by the ‘minpack.lm’ R package.^20^ The values of the algorithm parameters used during lineshape fitting are available in Supporting Information. To avoid falling into local minima, the fitting optimization is iterated a number of times proportional to the spectrum lineshape complexity, with signal parameter starting estimates that are randomly initialized for each iteration. After these iterations, the resolution with the least lineshape fitting error is chosen. After lineshape fitting, the areas below the signals are quantified, a specific fitting error for each signal is estimated (procedure explained in Supporting Information) and the signal parameter values are collected.

A graphical user interface (GUI) is used to select the metabolites to be profiled and the profiling method (area integration, signal deconvolution) for each of the signals. The GUI is also used to supervise the optimal value ranges for each chemical shift and half bandwidth to be used during lineshape fitting. In the case of chemical shift, the median range in both datasets was 0.006 ppm. In the case of half bandwidth, the median range was 50% of the median value. In the case of intensity, the tool automatically calculates the optimal value ranges by analysing the spectrum lineshape.

In the faecal extract dataset, 80 signals (66 through deconvolution and 14 through integration) from 52 different metabolites were profiled. In the serum dataset, 48 signals (43 fitted through deconvolution and 5 through integration) from 33 different metabolites were profiled. In addition, the signal parameter values and fitting errors were collected in both dataset profiling iterations.

### Prediction pipeline of expected signal parameter values

After profiling, the collected and outputted signal parameters were used to predict, in each signal, the expected spectrum-specific values (with their PIs) according to the information present in the other signals. These predictions can be used in future steps to evaluate the results and improve the fitting.

To make the spectrum-specific prediction of a signal parameter, the values of the parameter in the other signals are collected to create a dataset of predictors. Then, to enrich the quality of this dataset of predictors, three common steps in machine-learning processes are applied successively: data cleaning to minimize the influence of inaccurate values, feature selection, and feature engineering.^21^ After these enrichment steps, the signal parameter is predicted using the enriched dataset of predictors during the training of a random forest-based prediction model. The random forest algorithm is an ensemble learning method based on the bootstrap aggregation (also called *bagging*) of decision trees.^22,23^ The random forest algorithm solves the main drawback of bagging trees (the tendency to create similar decision trees with highly correlated predictions) by adding randomness to the tree construction process. Random forest models the possible nonlinear factors and showed higher performance during exploratory data analysis and lower variance during prediction. In addition, 0.632 bootstrap resampling is applied to minimize overfitting.^24^ Then, for each spectrum, the distribution which best represents the predictions generated during the bootstrap (see Figure 1 (c)) is estimated. From this distribution, the median value (with 95% PIs) is outputted as the spectrum-specific predicted value in the signal parameter analysed (Figure 1 (d)). The complete details of the prediction pipeline as well as the specifications regarding intensity prediction are available in Supporting Information.

It was considered that, if the predictions of parameters were not spectrum-specific, the best possible prediction of this parameter would consist of the median value found for this parameter in all spectra, having as 95% PIs the 95% central distribution of values. Accordingly, to evaluate the narrowness achieved in the spectrum-specific predictions generated, for each signal and parameter, the ranges of the 95% PIs of the spectrum-specific and the spectrum-unspecific predictions were compared.

In addition, a quality indicator based on the difference between the predicted signal parameters and the parameters obtained during profiling was calculated. For each one of the signal parameters with available information, the absolute difference was normalized to 0-1. Subsequently, the values obtained for each signal of each spectrum were averaged. As a result, a 0-1 ‘anomaly score’ was generated, which parameterizes how anomalous the signal parameter values obtained during profiling are.

### Evaluation of improvement in profiling data quality

The presence of both MS and NMR data made it possible to parameterize the improvement in the quality of profiling data. In both platforms, the concentration of 15 metabolites in the faecal extract dataset and 11 metabolites in the serum dataset was determined. Improvements in profiling quality in NMR data should be associated with an increase in Spearman’s rank correlation between the metabolite concentrations collected in NMR data and the ones collected in MS data. This indicator of profiling data quality was used to evaluate the profiling improvement after a new profiling iteration had been performed using the data of the predicted signal parameters. If the narrow and accurate PIs of signal parameters are used as parameter value ranges during fitting, more accurate resolutions during lineshape fitting should be expected.

In addition, the fitting error and the anomaly score were compared as quality indicators. To make this comparison, the worst quantification from the first profiling iteration according to the quality indicator was identified and corrected by its equivalent in the new profiling iteration. Then, the mean Spearman’s rank correlation between MS and NMR data was recalculated and the next worst quantification was identified. This process was iterated until all quantifications from the first profiling iteration had been corrected. It was expected that the better the quality indicator was the more able it would be to identify the quantifications to be corrected, so fewer corrections would be required to meaningfully increase the MS/NMR correlation.

## Results

### Accurate predicted values with narrow PIs which can be used to maximize profiling performance

The predictions generated (like the one in Figure 1 (d)) showed narrow spectrum-specific PIs for all the signal parameters analysed. For chemical shift, the median range in the spectrum-specific 95% PIs calculated in the faecal extract dataset was 4.7e-04 ppm. This value is lower than the bucket width (6e-04 ppm) and is a reduction of 75.8% in the median range in the spectrum-unspecific 95% PIs (1.9e-03 ppm) (Figure 2, top left). In the serum dataset, the median range in the spectrum-specific 95% PIs calculated was 1.9e-04 ppm, a reduction of 87.1% in the median range in the spectrum-unspecific 95% PIs (1.4e-03 ppm) (Figure 2, bottom left).

**Figure 2.**
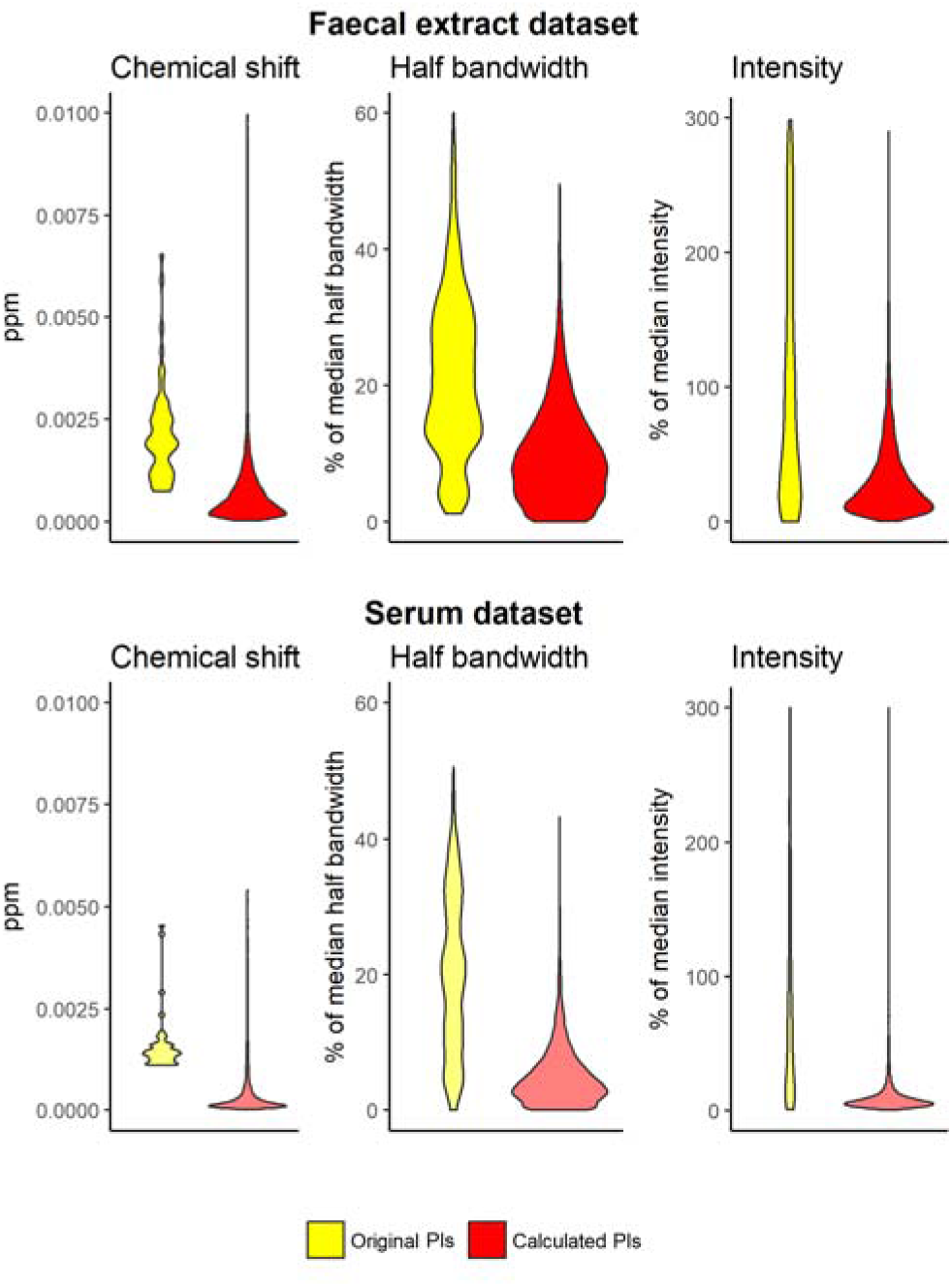
The spectrum-specific 95% PIs of the parameter values PIs are much narrower than the spectrum-unspecific 95% PIs. Chemical shift PIs are generally lower than the bucketing applied (6e-4 ppm). The narrow PIs enhance the performance of error minimization algorithms to end in the right local minimum.

For half bandwidth, the median range in the spectrum-specific 95% PIs calculated in the faecal extract dataset was 8.6% of the predicted half bandwidth. This value is a reduction of 58.4% in the median range in the spectrum-unspecific 95% PIs (20.6% of the predicted half bandwidth) (Figure 2, top middle). In the serum dataset, the median range in the spectrum-specific 95% PIs calculated was 4.0% of the predicted half bandwidth, a reduction of 80.3% in the median range in the spectrum-unspecific 95% PIs (20.1% of the predicted half bandwidth) (Figure 2, bottom middle).

For intensity, the median range in the spectrum-specific 95% PIs calculated in the faecal extract dataset was 22.2% of the predicted intensity. This value is a reduction of 92.8% in the median range in the spectrum-unspecific 95% PIs (309.9% of the predicted intensity) (Figure 2, top right). In the serum dataset, the median range in the spectrum-specific 95% PIs calculated was 6.9% of the predicted intensity, a reduction of 93.3% in the median range in the spectrum-unspecific 95% PIs (102.9% of the predicted intensity) (Figure 2, bottom right).

Apart from showing narrow PIs, the predictions also helped maximize profiling performance when they were used in a new profiling iteration. When all quantifications were corrected with the predicted information to improve the quality of the lineshape fitting, mean Spearman’s rho between MS and NMR metabolite concentrations increased 0.024 points (from 0.706 to 0.730) in the faecal extract dataset (Table 1; top) and 0.035 points (from 0.672 to 0.707) in the serum dataset (Table 1; down). Of the 25 correlations analysed, 21 of them increased their rho values, the maximum increase being 0.136 points and the maximum decrease 0.038 points. Rho improvements were especially important in the metabolites with the lowest correlation between the quantifications of both platforms: in the faecal extract dataset, the lowest rho value increased from 0.547 to 0.632; in the serum dataset, the lowest rho value increased from 0.515 to 0.589.

**Table 1.** The predicted signal parameter information increases Spearman’s rho correlation between metabolite concentrations in MS and NMR data in both datasets. There is a consistent increase in this profiling quality indicator when a new profiling iteration is performed using the PIs as new value ranges during lineshape fitting. The increase is most significant inthe metabolites whose profiling was most complicated in the original profiling iteration.

| Metabolites in faecal extract | Original Spearman's rho correlation | Spearman's rho correlation after new profiling iteration |
| --- | --- | --- |
| L-Isoleucine | 0.513 | 0.671 |
| D-Glucose | 0.556 | 0.589 |
| Glycine | 0.576 | 0.605 |
| L-Phenylalanine | 0.585 | 0.593 |
| L-Leucine | 0.659 | 0.690 |
| L-Valine | 0.672 | 0.699 |
| Citric acid | 0.690 | 0.695 |
| L-Tyrosine | 0.698 | 0.722 |
| L-Alanine | 0.726 | 0.737 |
| 3-Hydroxybutyric acid | 0.846 | 0.872 |
| Lactic acid | 0.871 | 0.901 |
| Metabolites in serum | Original Spearman's rho correlation | Spearman's rho correlation after new profiling iteration |
| Glycine | 0.547 | 0.658 |
| Hexanoic acid | 0.556 | 0.677 |
| 5-aminovaleric acid | 0.608 | 0.681 |
| L-Isoleucine | 0.629 | 0.632 |
| L-Alanine | 0.656 | 0.662 |
| L-Valine | 0.664 | 0.655 |
| L-Leucine | 0.697 | 0.715 |
| 2,4-Dihydroxypyrimidine | 0.718 | 0.744 |
| Succinic acid | 0.735 | 0.730 |
| Nicotinic acid | 0.765 | 0.784 |
| Phenylacetic acid | 0.794 | 0.799 |
| Glycerol | 0.809 | 0.771 |
| 3-Phenylpropionic acid | 0.844 | 0.873 |
| Lactic acid | 0.862 | 0.832 |

### High accuracy of the calculated anomaly score for detecting improvable quantifications

To parameterize the performance of the fitting error and the calculated anomaly score as quality indicators of quantification, the quality of quantifications was ranked using these quality indicators. This ranking was then used to gradually replace the worst ranked quantification in each metabolite by the equivalent one obtained in the new higher-quality profiling iteration. It was expected that this gradual replacement of quantifications would improve Spearman’s rank correlation between MS and NMR data with a logarithmic-like trend, as the first improved quantifications would provide the highest increases in the MS/NMR correlation.

In both datasets, the anomaly score as a quality indicator showed a logarithmic shape (Figure 3). Increase in the MS/NMR data correlation stopped improving after correcting approximately the 50% of the quantifications with worst anomaly score. Therefore, the anomaly score showed effectiveness at ranking the quantifications which might be further optimized. In comparison with the anomaly score, the fitting error showed a general lower effectiveness to detect improvable quantifications (as shown by the less logarithmic trend - Figure 3-).

**Figure 3.**
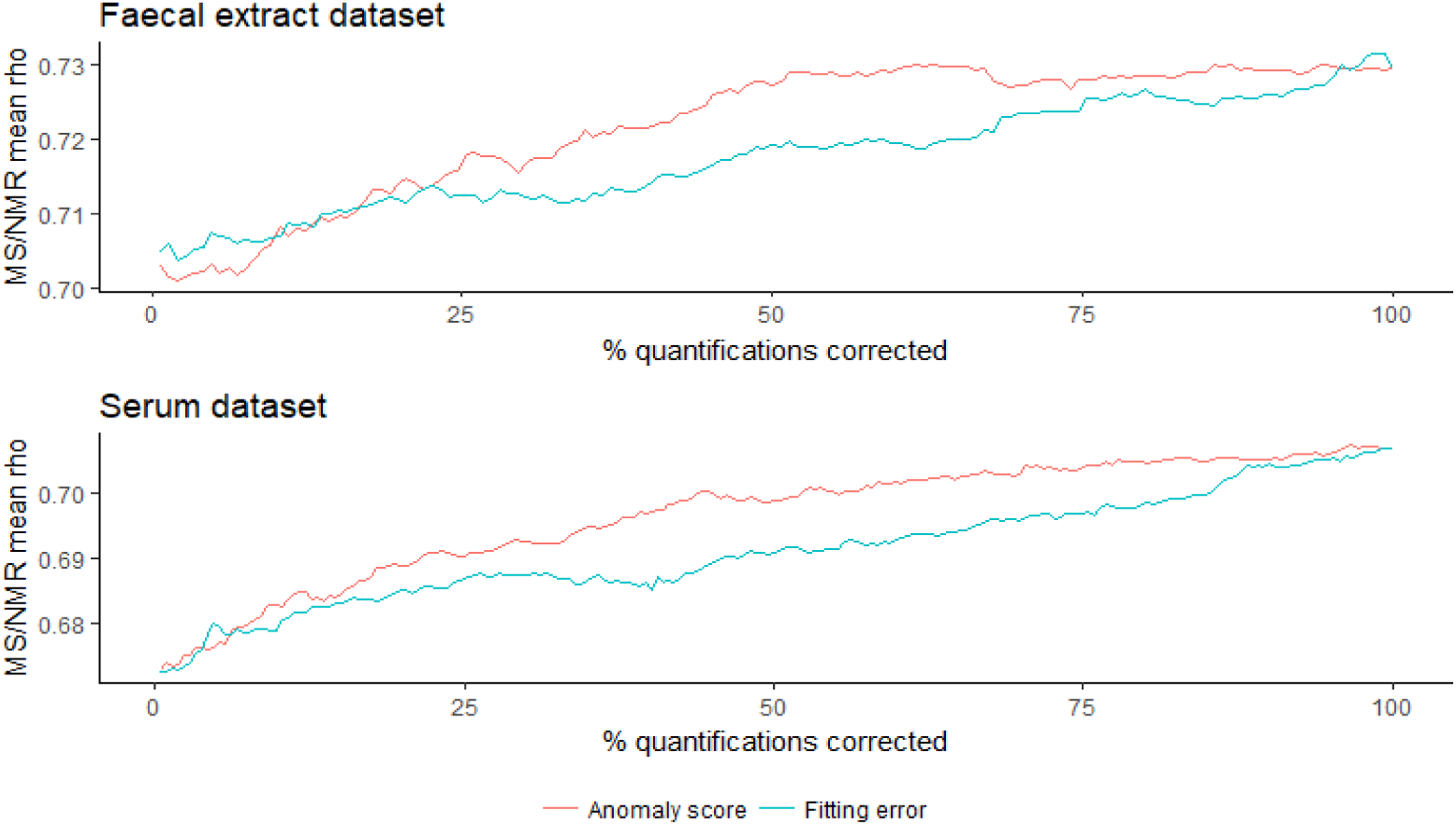
The calculated anomaly score helped identify quantifications which might be further optimized. In both datasets, the anomaly score showed higher performance than the fitting error ranking the quantifications which, if further improved, might further enhance the MS/NMR correlation.

The only subset of quantifications in which the fitting error performed better than the anomaly score was the worst quantifications in the faecal extract dataset (Figure 3; top). This seems consistent with the high importance of the intensity information in the fitting error compared to half bandwidth or chemical shift information. In the faecal extract dataset, high coefficient of variation and possible fitting of adjacent signals are challenges. Accordingly, occasional high distortions of estimated intensity can be found which are better parameterized by the fitting error. However, after detecting these extreme suboptimal quantifications, the fitting error would be less able than the anomaly score to find quantifications where the characterization of the signal does not behave as expected.

## Discussion

The results of the study showed that predicting signal parameter values with the information collected during a first profiling iteration helps maximize profiling performance. The improvement shown in this study has been demonstrated in biologically complex matrices and not in spike-in samples which cannot fully reproduce the usual complexity of metabolomics studies. Our study also presents a new quality indicator based on the information generated by our machine-learning based pipeline. This new quality indicator, called the anomaly score, may provide higher-quality information to improve the detection of suboptimal quantifications and enable the detection of wrong annotations, two current bottlenecks in metabolomic studies which contribute to the introduction of false positives and negatives into the metabolomics literature.^12,15,16^ In addition, our machine-learning based pipeline (contained in the ‘signparpred’ function in the ‘rDolphin’ R package) can be exported to any other profiling tool in any other programming language. The great benefits of our approach are mediated by the generation of predictions specific to each signal and each spectrum with accurate and narrow PIs. These high-quality predictions ensure that the algorithmic minimization of the signal fitting error prevents the pervasive problem of falling into wrong local minima when numerous parameter values are optimized (dozens of parameters in the case of complex lineshape fittings). Other approaches try to minimize this problem by creating narrow value ranges prior to profiling. However, when dealing with complex matrices, they may have limitations such as:

- Strict sample preparation or spectrum acquisition: difficulty of changing established protocols in labs, less flexibility to adapt the spectrum acquisition process to the properties of samples.
- Half bandwidth and chemical shift prediction: broadening of TSP signal mediated by protein, nonlinear patterns in certain signals in complex matrices, inability to handle unidentified metabolites.^10^
- Simultaneous lineshape fitting of all the signals of a same metabolite: variability in the relative intensity of signals depending on the matrix, challenges when signal chemical shift is not predicted exactly, inability to handle unidentified metabolites.
- Algorithm-based signal alignment: signal distortion, wrong annotations.^25,26^

In contrast, our approach is not dependent on restrictions or extensive previous information about signal properties: it only needs a flexible first profiling iteration that collects information for accurately characterizing the properties of the metabolite signals profiled and of the sample analysed. So, our approach provides a solution to the limitations listed above. Besides, the information obtained about the signal parameters of unidentified metabolites can be studied to find annotated signals with similar patterns (and consequently create valuable inferences about their structure and properties).

The maximization of the profiling quality shown in the results was not associated with a correlated decrease in the signal fitting error (the standard quality indicator outputted by NMR profiling tools). The mean fitting error of quantifications increased 0.26% in the faecal extract dataset and decreased 0.02% in the serum dataset. This suggests a ceiling in the performance of lineshape fitting approaches when matrices are complex. For example, they may give little importance to the lower intensity signals in the region analysed or not fully monitor the high-intensity baseline present e.g. in serum. Fitting information parameterizes not metabolite properties but spectrum properties. In contrast, the information generated with our prediction pipeline parameterizes metabolite properties. As a result, the new information generated by this workflow leads to next-generation quality indicators which are able to e.g. monitor wrong annotations because the associated chemical shift signal is not consistent with the information present in the whole dataset. This kind of quality control has the potential to filter out suboptimal quantifications more effectively. Consequently, it may be possible to profile many more metabolites without decreasing the profiling data quality.

The variability of chemical shift is one of the biggest challenges to progress in the automatic profiling of NMR datasets, and the PIs achieved during prediction tend to be even lower than the bucket width chosen. Thanks to this accurate chemical shift prediction, signals can be correctly assigned and the lineshape fitting performance maximized. The fact that chemical shift can be accurately predicted in faecal extract, a matrix with considerable variability in chemical shift and signal overlap, suggests that accurately predicting chemical shift in human urine is achievable. This matrix is of great interest to metabolomics. However, its complexity makes robust automatic profiling a real challenge, and it is recommended that some tools are not used in this matrix. A promising technique for maximizing the quality of NMR profiling in human urine through chemical shift prediction has recently been published.^13^ Nonetheless, this technique cannot be exported to NMR profiling tools because of licensing restrictions and it requires strict sample preparation and spectrum acquisition criteria. The machine learning pipeline we propose, when tuned to the special conditions of human urine and validated by comparison with MS data, may be a generalizable solution to the signal misalignment problem in human urine. In Supporting Information, we show the current results in a human urine dataset (not validated through MS data).

### Future directions

The benefits of our approach should also be observed in 2D NMR spectra and it may help solve some of their current limitations. Current use of 2D for quantitative purposes can be hindered by the lower proportionality between signal volumes and metabolite concentrations.^27^ This lower proportionality is mediated by the much higher complexity of the pulses used during spectrum acquisition and by the requirement of long experiment times which may lead to greater noise in the acquired data.^8^ Prediction of the signal properties may help increase this proportionality and expand its quantitative potential. It is plausible the workflow performed could also be helpful to solve the challenges observed in the profiling of datasets of other platforms such as MS. In MS, there are certain biological and technical factors that can interfere with the signal parameter values.^29^ At present there is considerable interest in solving the challenges present in these datasets. The collection of signal parameter values and the use of our approach may help to this purpose.

## Conclusions

Until now, profiling tools required restrictions (e.g., in the analysed matrix, the preparation of samples, or the acquisition of spectra) to ensure that their workflows could be generalizable to the analysed dataset. The results of this study showed that profiling workflows can be generalizable to any kind of dataset if the information necessary to ensure this generalization is previously collected from the same dataset to be profiled. The multicollinearity present in this collected information enables a narrow prediction of the expected signal parameters which is robust to the noise present in this information. As a result, the quality of the signal parameter values achieved during profiling can be maximized and, therefore, the quality of automatic metabolite profiling can be maximized. Similar achievements might be followed if trying to implement similar solutions in other metabolomics challenges where there are factors provoking a variability in the signal parameters which needs to be monitored.

## Notes

The authors declare no competing financial interest.

## Acknowledgments

We thank Dr. Oscar Yanes from URV/CIBERDEM for helpful discussions, and Dr. Roger Paredes and the Microbial Genomics group from IrsiCaixa for giving us access to NMR spectra of fecal samples.

